# Effects of maternal deprivation and complex housing on pro-social behavior in rats: An automated, operant task examining motivation to liberate a trapped conspecific

**DOI:** 10.1101/2020.11.30.403386

**Authors:** Aikaterini Kalamari, Jiska Kentrop, Chiara Hinna Danesi, Evelien J.M. Graat, Marinus H. van IJzendoorn, Marian J. Bakermans-Kranenburg, Marian Joëls, Rixt van der Veen

## Abstract

Early life environment influences the development of various aspects of social behavior, particularly during sensitive developmental periods. Here, we aimed to study how challenges in the early postnatal period or (early) adolescence affect pro-social behavior. To this end, we adapted an existing paw operated liberation task to an automated operant task, to measure motivation (by progressively increasing required lever pressing) to liberate a trapped conspecific. Liberation of the trapped rat resulted either in social contact, or in liberation into a separate compartment. Additionally, a condition was tested in which both rats could freely move in two separate compartments and lever pressing resulted in social contact. When partners were not trapped, rats were more motivated to press the lever for opening the door than in either of the trapped configurations. Contrary to our expectation, the trapped configuration resulted in a *reduced* motivation to act. Early postnatal stress (24h maternal deprivation on postnatal day 3) did not affect behavior in the liberation task. However, rearing rats from early adolescence onwards in complex housing conditions (Marlau cages) reduced the motivation to door opening, both in the trapped and freely moving conditions, while motivation for a sucrose reward was not affected.

## Introduction

Emotional contagion, i.e. the capacity to experience and share the emotions of others, is considered an evolutionary well-preserved mechanism that helps individuals to survive, not only in a dangerous savanna, but also within social groups (Preston and De Waal, 2001; de Waal, 2008; Kim et al., 2019). Rodents, like many other animals, experience and learn from emotional contagion, as shown in studies on emotional contagion for pain (Church, 1959; Langford, 2006; Langford et al., 2010; Atsak et al., 2011; Pereira et al., 2012; Li et al., 2014; Cruz et al., 2020) and observational fear conditioning (Kavaliers et al., 2003; Jeon et al., 2010; Kim et al., 2010; Allsop et al., 2018; Keum and Shin, 2019; Nomura et al., 2019).

Acting pro-socially, that is, showing behavior that benefits others, builds upon emotional contagion, but additionally requires perspective taking and empathic concern about the other’s state (de Waal, 2008; Chen, 2018; Sivaselvachandran et al., 2018). Initially, empathic concern and perspective taking were thought to be characteristics unique to humans, but an increasing number of experimental studies challenge this idea and suggest that pro-social behavior and its underlying mechanisms can also be studied in non-human primates and other animals (De Waal and Preston, 2017; Pérez-Manrique and Gomila, 2018; Sivaselvachandran et al., 2018). This also holds true for rodents. Thus, some studies have shown that rats help distressed conspecifics in laboratory settings, e.g. pressing a lever to lower a rat that is suspended in the air (Rice and Gainer, 1962) and allowing a soaked rat to escape a pool of water (Sato et al., 2015). Bartal and colleagues (2011) developed a model in which a rat was trapped in a cylinder that could be manually opened by a conspecific. After being trained to operate the cylinder door, a high percentage of rats actively liberated trapped familiar conspecifics. Rats did not open the cylinder when it was empty or contained a toy rat, suggesting that the liberation act has reinforcing properties. In a follow-up study, rats were shown to liberate familiar but not unfamiliar conspecifics (Bartal et al., 2014), demonstrating that previous social experiences shape helping behavior later in life, in line with other studies (Langford et al., 2010; Burkett et al., 2016; Rogers-Carter et al., 2018).

We adapted this pro-social liberation model to an operant set-up where a rat is trapped in a cylinder blocked by automated doors. These doors can be opened by a free rat in an adjacent compartment, by means of lever pressing. To assess the motivation to liberate a trapped conspecific, the number of presses needed to open the doors is progressively increased over sessions, increasing the cost of liberation (in contrast to a single action). To distinguish between motivation to liberate a distressed conspecific and motivation for social contact (Silberberg et al., 2014), the task was performed using two configurations: one in which opening the door gave the trapped rat access to the cage of the liberator, enabling social contact; and one in which opening the door gave the trapped rat access to a separate compartment. Additionally, a third configuration was tested in which both rats could freely move in separate compartments and lever pressing resulted in gaining access to the other’s compartment.

Ultrasonic vocalization (USV) is an important communication channel for rats and might influence behavior in the pro-social liberation task. For adult rats, USVs can be divided in two categories that reflect different affective states: ‘alarm’ calls in the range of 18-32 kHz (referred to as 22KHz calls); and ‘appetitive’ calls in the range of 33-96 kHz (referred to as 50 KHz calls) (Portfors, 2007; Takahashi et al., 2010; Wright et al., 2010; Brudzynski, 2013; Wöhr and Schwarting, 2013). Calls in the 22 kHz domain are emitted during aversive situations such as direct danger, approaching danger or emotional distress, whereas 50 kHz calls are usually emitted in pleasant situations and function to establish and maintain contact with conspecifics (Simola and Brudzynski, 2018; Brudzynski, 2019). Social enrichment can lead to an increase in ultrasonic communication and approach behavior in response to appetitive calls (Brenes et al., 2016).

Social behavior is affected by previous experiences, especially during sensitive developmental periods such as the early postnatal period and (early) adolescence (Marco et al., 2011; Sandi and Haller, 2015; Tzanoulinou and Sandi, 2016). In rodents, early life adversity in the first two weeks after birth –by depriving pups of maternal care or providing pups with poor quality of maternal care– can negatively affect social behavior, although this has not been extensively studied yet (Bonapersona et al., 2019). Here we used 24h deprivation of maternal care on postnatal day 3, presumably modeling neglect or trauma. This model has previously shown to induce HPA-axis changes (Workel et al., 2001; Schmidt et al., 2004; Enthoven et al., 2010) and long-lasting structural (Loi et al., 2014; Sarabdjitsingh et al., 2017) and functional (Oomen et al., 2010, 2011; Derks et al., 2016; Loi et al., 2017) changes in the brain. We have shown that this deprivation negatively affects adult behavioral inhibition and social discrimination (Kentrop et al., 2016, 2018), leading to the hypothesis that pro-social behavior may also be impaired.

Though sensitive periods are mainly studied in the context of early life adversity and its negative consequences for brain development, it is likely that a favorable early life environment might benefit development. For example, communal nesting, in which two or more mothers raise their pups together in a shared nest, was found to enrich the repertoire of social behaviors in mice (Branchi and Alleva, 2006; Branchi and Cirulli, 2014). Later in life, the exposure to a more naturalistic and ‘enriched’ environment, comprised of both social and physical enrichment and regular exposure to novelty increases brain plasticity and may positively influence the development of social skills (Würbel, 2001; Gubert and Hannan, 2019). Like in humans, the period of adolescence is a period in rodents in which brain circuitry implied in social behavior is still in development (Fuhrmann et al., 2015; Casey et al., 2019). Studies applying this ‘enrichment’ during the adolescence period report reduced anxiety and enhanced learning, memory, and social behavior later on (van Praag et al., 2000; Simpson and Kelly, 2011; Crofton et al., 2015). Here, we housed animals from adolescence onwards in large cages that contain all aspects of an ‘enriched’ lab environment and which we refer to as complex housing. We have recently found differential effects on social play in adolescence and social interest in adulthood in animals housed in these complex cages compared to standard housed animals (Kentrop et al., 2018), which might also be reflected in pro-social behavior.

I **Error! Bookmark not defined.**n summary, two separate experiments were conducted to investigate the influence of (early) life experiences on pro-social behavior in adult rats: 1) manipulation of the early postnatal environment through 24h MD on postnatal day 3 and 2) manipulation of the adolescent environment through complex housing from postnatal day 26 onwards, up and throughout testing in adulthood. All rats were tested in our automated operant version of the liberation task in adulthood.

## Materials and methods

### Animals

Male and female Wistar breeding rats were obtained from Charles River Laboratories (Arbresle, France) at 8-10 weeks of age. All experimental rats were bred in-house, except for 7 pairs of males from pilot experiment B that originated directly from Charles River. Only male rats were included in the experiments. Standard housed rats were kept in type IV Makrolon cages (37 × 20 × 18 cm) in temperature (21°C) and humidity (55%) controlled rooms with a 12h light–dark cycle (lights on from 08:00-20:00) during breeding, and a reversed cycle (lights on from 20:00-08:00) from postnatal day 21 onwards. Unless stated otherwise, food and water were available ad libitum and all cages were provided with a woodblock as standard cage enrichment. Clean cages were provided and general health status was checked on a weekly basis. In the MD and complex housing experiments, animals were semi-randomly assigned to the experimental or control groups; with the constraint that a maximum of 2 rats from the same litter were assigned to the same group to minimize litter effects. Rats were tested in adulthood (90+ days of age) in pairs; one acting rat, referred to as *test rat*, and one stimulus rat, referred to as *partner*. Rats participated either as test rat or partner. Where possible, siblings were housed together and became partners. All rats were previously tested in a pro-social 2-choice task (Kentrop et al., 2020), with at least 2 weeks rest before entering the current experiment; the test rats in the current study were partners in the pro-social 2-choice task and vice versa. The present study consists of a series of pilot experiments, an early life stress experiment with maternal deprivation (MD); and a complex housing experiment, see Figure 1A for a schematic overview of all experiments. Experiments were approved by the Central Authority for Scientific Procedures on Animals in the Netherlands (CCD project AVD115002016644). Animal care was conducted in accordance with the EC Council Directive of November 1986 (86/609/EEC).

**Figure 1:**
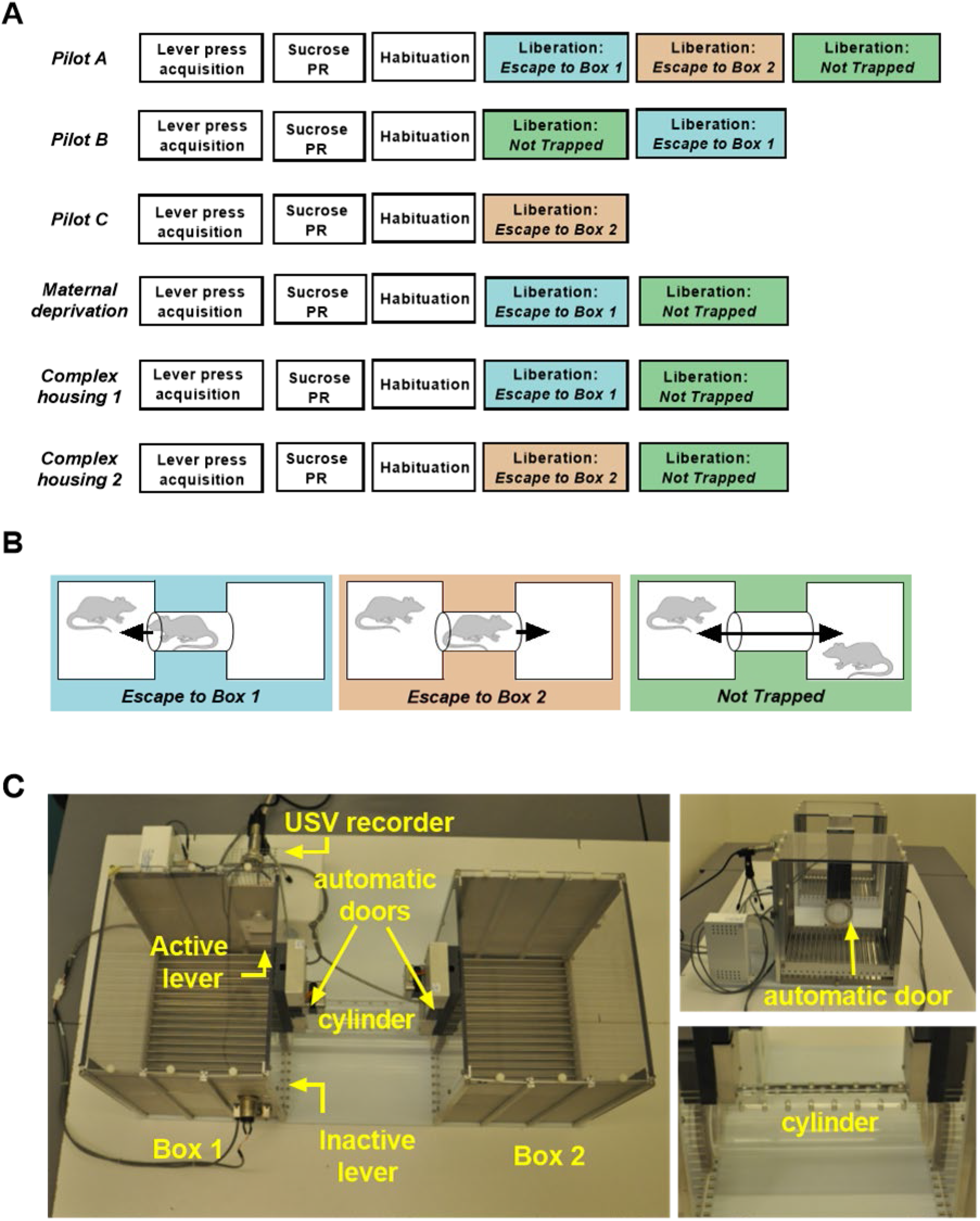
(A) Schematic overview of behavioral protocols used in the three pilots (A,B,C), the maternal deprivation experiment and the complex housing experiment. PR = progressive ratio. (B) Three different configurations in the liberation task were tested: 1) *Escape To Box 1* in which the partner was trapped in the cylinder and could be liberated from the cylinder into box 1 (the test rat compartment), 2) *Escape To Box 2* in which the partner was trapped in the cylinder and could be liberated into box 2, and 3) *Not Trapped* configuration in which the partner was situated in box 2 and lever presses of the test rat opened both doors. (C) Liberation task set-up: 2 operant chambers are connected through a removable cylinder. The cylinder can be closed on both sides by automated mechanical doors made of transparent Plexiglas with holes that allowed rats to see, smell and hear, but not touch each other.

### Breeding and maternal deprivation

Breeding started after the rats had been familiarized with our animal facility for at least 2 weeks. Two females were paired with 1 male for 10 days. After separation from the male, females stayed together for another week and were then individually housed to prepare for birth. Paper towels were provided to the mothers as nesting material. The day of birth was considered postnatal day 0. On postnatal day 3 the sex of the pups was determined and when necessary litters were culled to a maximum of 10 or supplemented to a minimum of 6 by adding pups from surplus animals from culled litters. Mothers and pups were placed back into their home cage within 2 min, except for litters in the MD experiment. During deprivation, the pups stayed together in their home cage (without the dam, which remained single housed in a separate cage without pups, with food and water ad libitum) and were transported to a different room. The cage with pups was placed on a heating pad (33°C) to prevent hypothermia; the pups were left undisturbed and they were not fed. After 24h the cage was taken back to the original room and the mother was reunited with her litter. The week following deprivation, mothers from deprived litters were not provided with additional nesting material.

### Weaning and complex housing

Pups were weaned on postnatal day 21. Pups from the pilot experiments, MD experiment and the standard housing condition in the housing experiment, were pair-housed in type IV Makrolon cages after weaning. Rats from the housing experiment assigned to the complex housing condition were housed in type IV Makrolon cages with 3 to 4 males from postnatal day 21 to 26 and then transferred to Marlau^™^ cages (Viewpoint, Lyon, France), housing 10 males per cage. Marlau cages are large, enriched cages (60 × 80 × 51 cm) that have 2 floors and provide a complex and challenging environment for the rats (Fares et al., 2013). The first floor contains a big compartment with three running wheels, a shelter, ad libitum access to water, 2 woodblocks, and a climbing ladder to the second floor, where a maze has to be passed to gain access to a tube leading to the food compartment on the first floor. Via a one-way passage rats can regain access to the bigger first floor compartment. The maze was changed once per week (alternating between 12 different configurations), assuring novelty and sustained cognitive stimulation. Territorial dominance was avoided by the presence of 2 openings on each side of the maze. A more detailed description of the experimental setup is given elsewhere (van der Veen et al., 2015).

### Boldness test

To determine which rats would become test rat and partner, rats were subjected to a boldness test. Boldness was measured by opening the lid of a cage half-way and measuring for each rat how much time was needed to rear and place at least 1 paw on the edge of the cage. This test was repeated 3 times on separate occasions and rats were ranked based on their cage emergence latency; the boldest rats (i.e. with the shortest emergence latency) were selected as test rats and the others as partners. The selection based on boldness was made to ensure that none of the test rats would have to be excluded from the experiment due to non-performance as a result of fear (replicated from the protocol by Bartal et al., 2011). For standard housed rats, the boldness test was conducted in the home cage and the selection was made within each pair. Complex housed rats were first removed from their complex home cage into type IV Makrolon transport cages (to which they were extensively habituated) and were left undisturbed for 30 min. After 30 min, rats were transferred one by one to an empty transport cage and tested individually. Rats with the highest boldness rank were selected as test rats.

### Acquisition of lever pressing and progressive ratio for sucrose

#### Temporary mild food restriction and sucrose pellet habituation

In order to acquire lever pressing before the start of the liberation task, rats were trained with sucrose pellets in separate operant cages. During acquisition of lever pressing, animals were mildly food restricted (3-4.5g chow/100g body weight) to attain 90-95% of their free fed bodyweight. During the periods of food restriction, animals were weighed twice per week to monitor body weight. The week before training rats received sucrose pellets in the home cage on three occasions to habituate to the taste of the pellets.

#### Apparatus

Acquisition of lever pressing took place in separate operant chambers (30.5 × 24.1 × 21 cm, Med Associates, St. Albans, VT, USA) equipped with 2 retractable levers, a cue light above each lever, a house light (which illuminated the chamber with dim light) and a pellet magazine in between levers that provided rats with a sucrose pellet (45 mg, Formula P; Bio-Serv). Rats could earn a sucrose pellet by pressing the active lever until the fixed ratio (FR) criteria was reached. Presses on the inactive lever were registered but had no consequences. The operant chambers were enclosed in larger boxes equipped with exhaust fans that assured air renewal and masked background noises. Experimental contingencies were controlled and data were collected using MED-PC version 14.0 (Med Associates).

#### Behavioral procedure

##### Habituation

Rats were habituated to the operant chambers in a 15 min session in which every minute the cue light above the active lever was turned on for 20 sec in combination with delivery of a sucrose pellet.

##### Acquisition training

A training session lasted 30 min and rats were trained twice a day (morning and afternoon). Training started with a fixed ratio 1 (FR1) schedule of reinforcement where 1 active lever press was required for the delivery of a sucrose pellet. Initially, rats were trained on an FR1 protocol in which both levers were active and trials were separated by a 10 sec inter-trial-interval (ITI) until the acquisition criteria of 30 pellets within a session was reached. Rats were subsequently trained on an FR1 (with an acquisition criteria of 30 pellets), FR3 (30 pellets) and FR5 (18 pellets) protocol with 20 sec ITI, with both the active and inactive lever present. Upon reaching the required number of presses to complete the fixed ratio, all levers retracted, the cue light above the active lever turned on to signal reward delivery and a sucrose pellet was dispensed in the pellet magazine.

##### Progressive ratio

To assess the motivation to work for a food reward, a progressive ratio (PR) protocol was conducted in which the number of required presses to earn a sucrose pellet progressively increased from 1 to 2, 4, 6, 9, 12, 15, 20, 25, 32, 40, 50, 62, 77, 95, 118, 145, 178, 219, 268, 328, etc. according to (Richardson and Roberts, 1996). A PR session lasted maximally 90 min or ended as soon as a rat did not reach the new ratio within 15 min of the previous one. PR was conducted twice, on 2 separate days. After the last PR test, rats returned to ad libitum feeding and were allowed to recover for at least 1 week before behavioral testing in the liberation task. For all acquisition training and PR sessions, the total numbers of rewards and active and inactive lever presses were recorded.

### Liberation task

#### Apparatus

The liberation task was conducted in a set-up consisting of 2 operant chambers (29.5 × 23.5 × 27.3 cm, Med Associates, St. Albans, VT, USA) placed 25 cm apart and connected by a removable transparent Plexiglas cylinder (25 cm in length and 7.5 cm in diameter) with holes that provided fresh air (6 mm in diameter) for rats that were trapped in the cylinder (Figure 1C). The cylinder could be closed on both sides by automated mechanical doors made of transparent Plexiglas with holes (6 mm in diameter) that allowed rats to see, smell, hear but not touch each other. Depending on the configuration tested, 1 or both doors were programmed to open in response to lever pressing by the test rat. Box 1, the test rat compartment (left side), contained 2 levers; 1 active lever with a cue light situated above it, and 1 inactive control lever. To record ultrasonic vocalizations a microphone was placed behind a piece of wire mesh integrated in the upper part of the wall in box 1. Experimental contingencies were controlled by MED-PC IV Version 4.2 software (Med Associates, St. Albans, VT, USA). During testing, the number of active and inactive lever presses, door openings, and door opening latencies were recorded. Experiments were conducted in red light conditions during the dark (active) phase.

#### Behavioral procedure

##### Habituation to box and levers

Rats were habituated to the set-up for 20 min without levers and with full access to box 1, box 2 and the cylinder (i.e. both doors were open). Following this, on 4 consecutive days rats were placed in the setup for 20 min on an FR1 (2x) and FR3 (2x) protocol in which both levers were present, the cylinder doors were closed and pressing the active lever resulted in door opening of both doors for 1 min, giving access to the cylinder and box 2. Rats coupled lever pressing in this cage to door opening, and active pressing quickly declined.

##### Configurations in liberation task

Three different configurations in the liberation task were tested (Figure 1B): 1) *Escape to Box 1* in which the partner was trapped in the cylinder and could be liberated from the cylinder into box 1 (the test rat compartment), 2) *Escape to Box 2* in which the partner was trapped in the cylinder and could be liberated into box 2 and 3) *Not Trapped* in which the partner was situated in box 2 and lever presses of the test rat opened both doors. In all configurations, test rats were placed in box 1 and could open the cylinder door(s) by lever pressing, thereby liberating their partner from the cylinder or gain access to each other.

##### (Adapted) Progressive Ratio

To assess the motivation to liberate a trapped cage mate, a progressive ratio protocol was introduced. As a proxy for motivation, the required number of presses needed to liberate the cage mate per ratio progressively increased over sessions from 1 to 3, 5, 7, 9, 12, 15, 20, 25, 32, 40, 50, 62, 77, etc. (adapted from Richardson and Roberts, 1996) until rats reached a breakpoint (Figure 2B). The breakpoint is the maximum amount of presses animals are willing to make within 10 min to liberate their partner. Thus, if a rat completed 5 ratios in the PR, this means that the maximum amount of presses was 9 and the sixth ratio was not completed (i.e. the rat did not reach 12 presses within 10 min). The difference with the PR for sucrose is that the ratios did not progress within a session, but between sessions. Rats were tested once a day in 1 or 2 consecutive sessions in which the test rat was placed in box 1 and could open the door(s) by reaching the required amount of lever presses within 10 min (see Figure 2A). As soon as rats completed a given ratio, the door(s) opened and the session ended 5 min later. If rats did not complete a ratio, the session ended after 10 min and both rats were taken out of the test. In order to progress, rats needed to complete a given ratio twice in maximally 3 sessions (Figure 2B).

**Figure 2:**
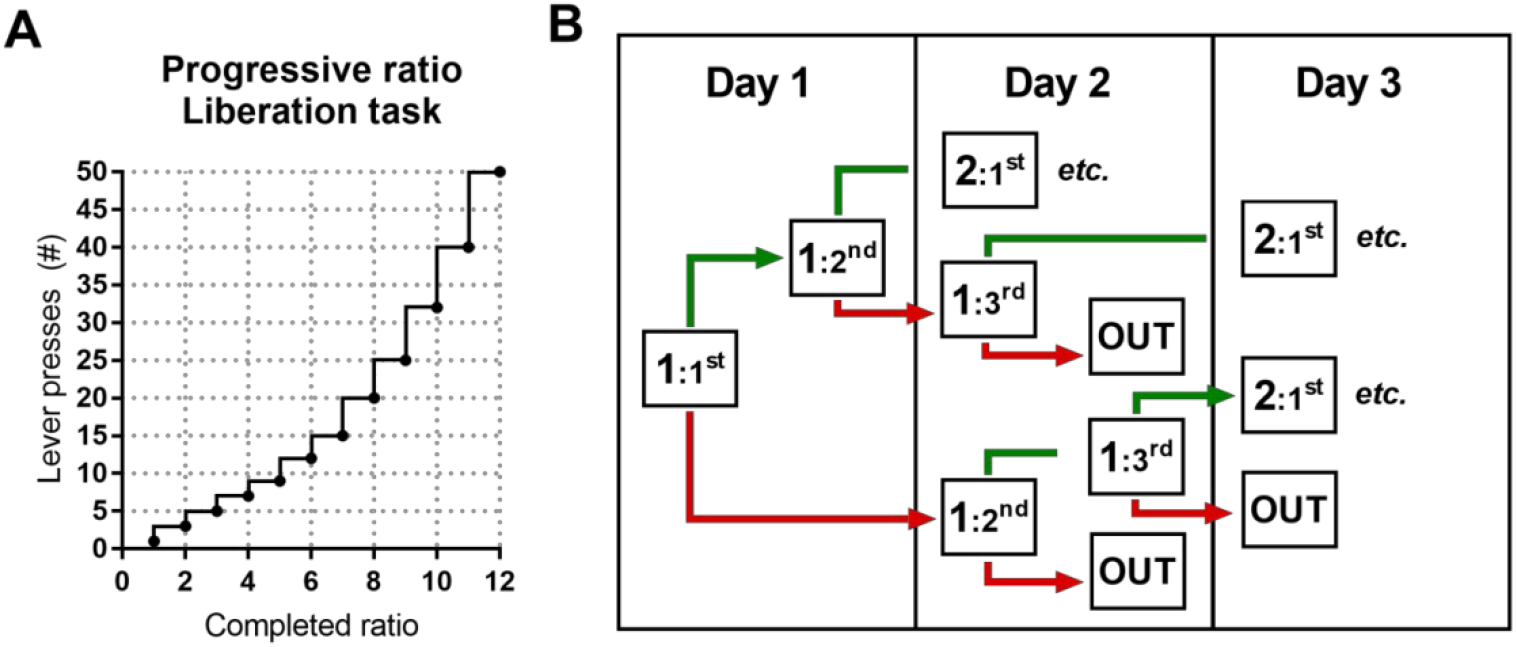
Schematic representation of the adapted progressive ratio protocol in the liberation task. (A) The progressive increase in number of required lever presses to complete a ratio and open the cylinder door(s) (adapted from Richardson & Roberts, 1996). (B) Decision tree in the liberation task. If during the first attempt on a given day rats reached the required number of lever presses (ratio completed; in green), a second attempt followed. If during the first attempt on a given day rats failed to reach the required number of lever presses (ratio not completed; in red), rats were tested again the next day. Rats needed to complete a given ratio twice in maximally 3 attempts. When they failed to do this, the preceding ratio was recorded as the maximum ratio reached.

### Recording and analysis of ultrasonic vocalizations

For the complex housing experiment, USVs of the first (FR1) session in the liberation task were analyzed. Ultrasonic vocalizations were recorded with Avisoft Bioacoustics RECORDER version 4.2.18 (Avisoft Bioacoustics, Berlin, Germany) and analyzed with UltraVox XT version 3.2.108 (Noldus Information Technology, Wageningen, the Netherlands). It was not possible to distinguish calls from individual rats, so USVs represent the communication between both rats. Spectograms were produced from the recordings by fast Fourier transformation (977 Hz frequency resolution, 1.024 ms time resolution, 256 FFT-length, 100% Frame, and 70% time window overlap). A lower cut-off-frequency was used to remove all background noise below 18 kHz. Calls were labeled using automated parameter measurements. Duration and start time of USVs were extracted for further analyses. Sounds were labeled as calls if the duration was at least 3 ms. Multiple fragments of calls (or syllables) were labeled as one call if the silence between fragments was less than 3 ms. Two categories of calls were distinguished: 22kHz alarm calls (in the range of 18-32kHz) and 50kHz appetitive calls (in the range of 32-96 kHz) according to (Wöhr and Schwarting, 2013). Calls with frequencies in both categories were not included in the analysis. The reported variable is the percentage of time that rats emit 22 or 50 kHz before and after door opening (calculated as (total duration of 22kHz or 50kHz calls (sec)/task duration before or after door opening (sec))*100%).

### Statistical analysis

Statistical analyses were performed using SPSS for windows version 23 (IBM, United States). Results are presented as mean ± SEM. No outlying data points (defined as 3.29 standard deviations below or above the mean) were detected. To compare emergence latency between test rat and partner rat in the boldness test, an independent samples Student’s *t*-test was conducted for the pilot or two way ANOVAs for the early life experiments with role (test- or partner rat) and treatment (no-MD vs MD; or standard housing vs complex housing) as between subjects factors. Performance in the progressive ratio task for sucrose was analyzed with independent Student’s *t*-tests in the MD and complex housing experiments. Behavior in the liberation task for pilot experiment A was analyzed using a repeated measures ANOVA with configuration (3 different configurations) as within subjects factor, followed by post-hoc *t*-tests with Bonferroni correction for multiple testing. A paired samples *t*-test was conducted to analyze the 2 different configurations in pilot B. Comparisons across pilot experiments were analyzed using independent Student’s *t*-tests. For the MD and complex housing experiments, (early) life effects were analyzed with independent Student’s *t*-tests for each configuration separately (since not all animals were tested in each configuration) and configuration effects were analyzed with repeated measures ANOVAs. To assess the relation between motivation to work for a sucrose reward and motivation to liberate a trapped cage mate, Pearson’s correlations were computed in the complex housing experiment.

To analyze the effect of housing on ultrasonic communication before door opening, independent Student’s *t*-tests were performed. To test the effect of door opening (liberation) on USVs, repeated measures ANOVAs were conducted within each call type with door opening (before vs after) as within subjects factor and housing as between factor, followed by post-hoc *t*-tests with Bonferroni correction for multiple testing if applicable. For a proper comparison of USVs before and after door opening, the pairs in which the test rat did not open the cylinder door(s) within the 10 min of the test (depicted in orange in Figure 6 and 7), were not included in this analysis. To compare the amount of alarm calls and appetitive calls in the liberation task, repeated measures ANOVAs were conducted with call type (22 kHz vs 50 kHz calls) and door opening (before vs after) as within-subjects factor and housing as between-subjects factor. To determine if ultrasonic communication before door opening (liberation) was related to the door opening latency, multiple regression analyses were performed for 22 kHz and 50 kHz calls separately with door opening latency as the dependent variable and USVs and housing condition as predictor variables. Where appropriate, partial eta squared *η_p_^2^*, Hedges’s g_s_ or g_av_ are used to present effect sizes according to (Lakens, 2013).

**Figure 6:**
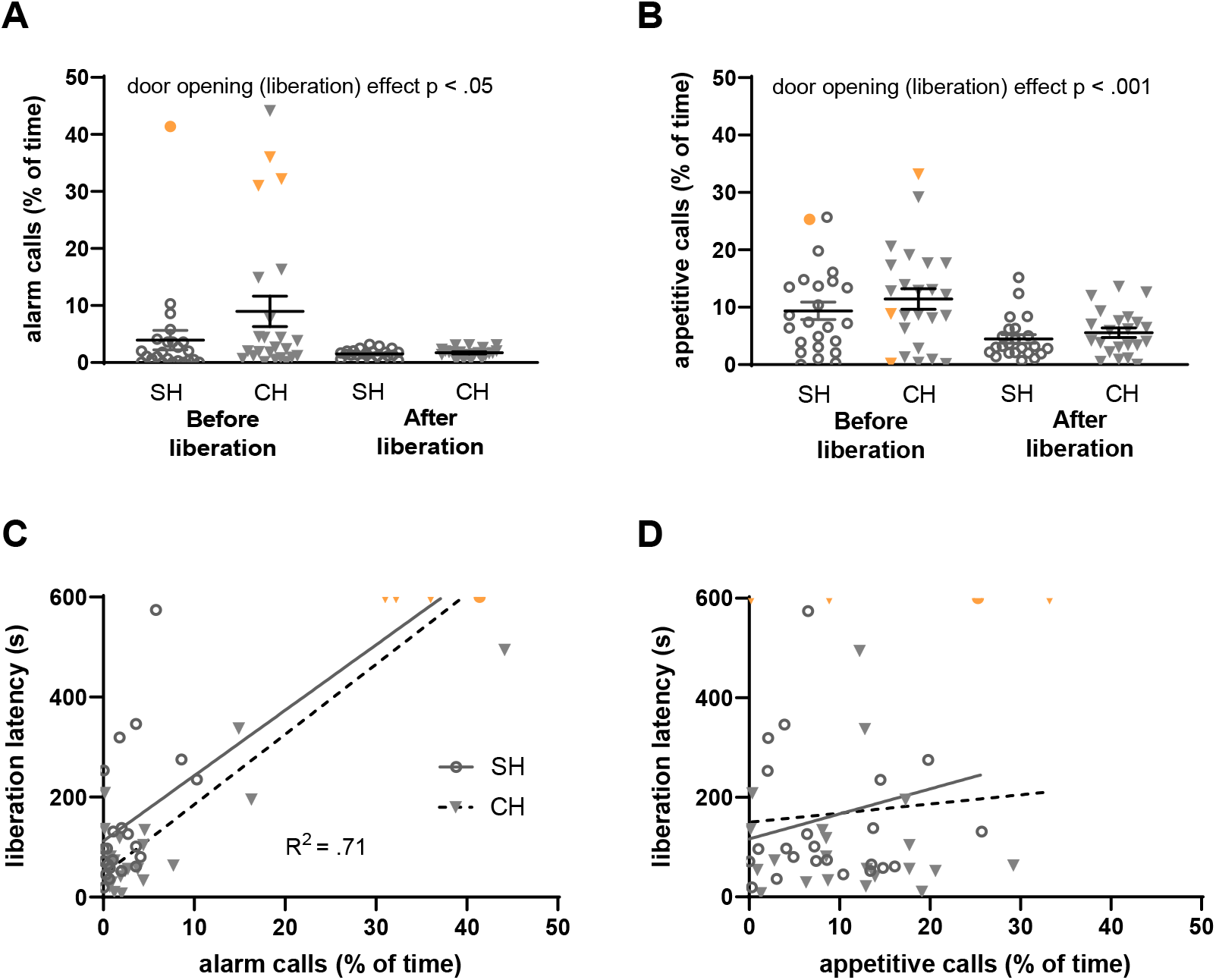
Ultrasonic vocalizations (USVs) recorded during the first (FR1) trial of the *Escape to Box 1* configuration, in the complex housing experiment. Vocalizations are presented as the percentage of time rats emit these USVs. Per experimental group, 24 pairs of rats were included of which 4 rats (3 complex housed and 1 standard housed rat) did not liberate their cage mates (data points depicted in orange). (A) 22 kHz (alarm) calls and (B) 50 kHz (appetitive) calls before and after liberation for standard and complex housed rats. (C) The relation between 22 kHz vocalizations (before liberation) and liberation latency and (D) the relation between 50 kHz vocalizations (before liberation) and liberation latency. SH: standard housed (black line), CH: complex housed (dashed line). Data represent individual data points and mean ± SEM.

**Figure 7:**
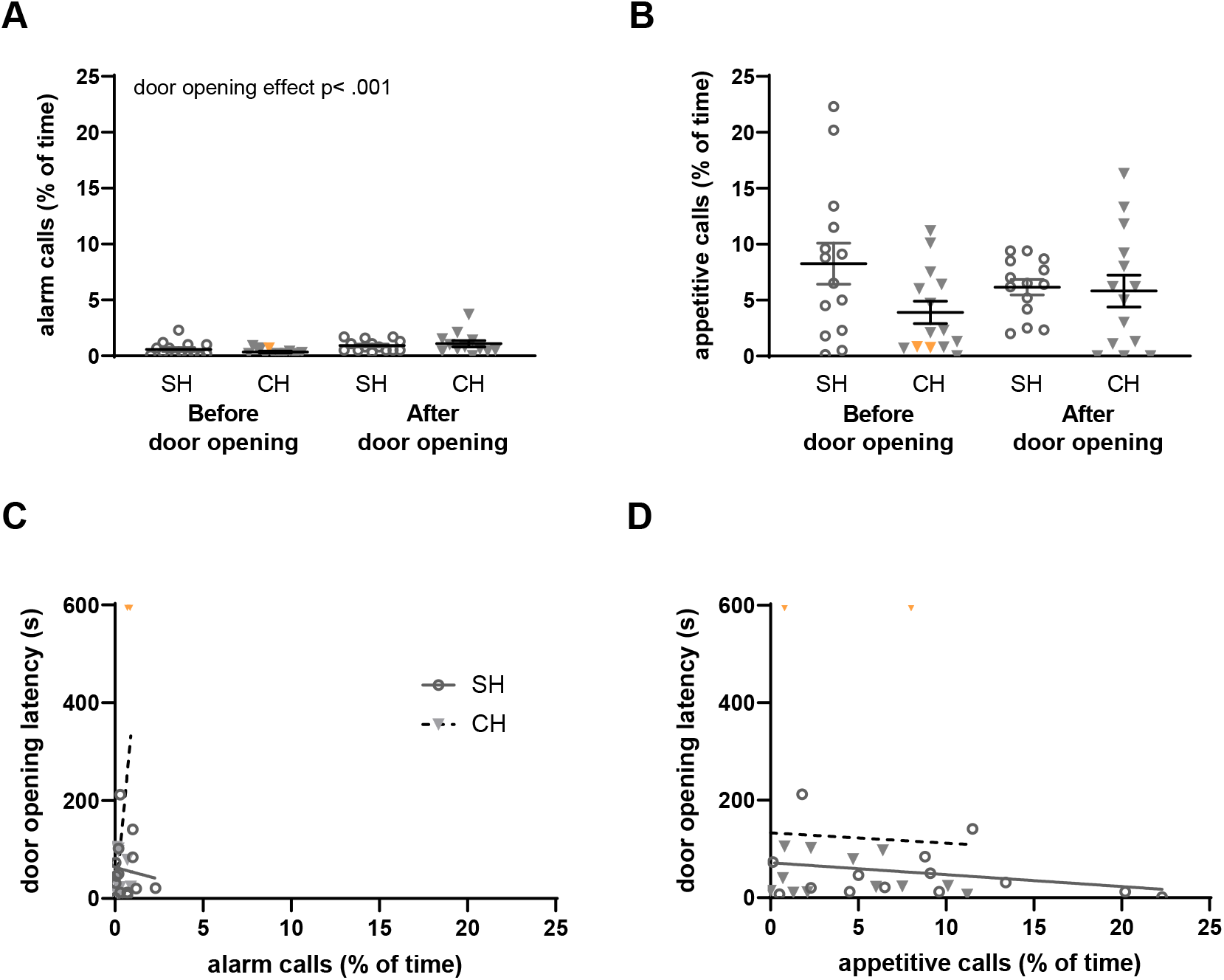
Ultrasonic vocalizations (USVs) recorded during the first (FR1) trial of the *Not Trapped* configuration in the complex housing experiment. Vocalizations are presented as the percentage of time rats emit these USVs. Per experimental group, 14 pairs of rats were included of which 2 rats (complex housed) did not open the doors (data points depicted in orange). (A) 22 kHz (alarm) calls and (B) 50 kHz (appetitive) calls before and after door opening for standard and complex housed rats. (C) The relation between 22 kHz vocalizations (before door opening) and door opening latency and (D) the relation between 50 kHz vocalizations (before door opening) and door opening latency. SH: standard housed (black line), CH: complex housed (dashed line). It should be noted that the amount of time animals emitted alarm calls was very low in this configuration. For this reason, the scaling of Figure 7 differs slightly from Figure 6. Data represent individual data points and mean ± SEM.

## Results

### Boldness measures

The most bold animal of a pair was selected to be the test rat in the liberation task according to Bartal et al. (2011). In the pilot experiments, the average emergence latency of designated test rats was not significantly different from the emergence latency of the designated partner rats (*t*(34) = −1.61, *p* = .12, g_s_ = 0.53, with test rats 30.3 sec ± 4.0 sec and partners 38.9 sec ± 3.6 sec). In the maternal deprivation and complex housed experiments, test rats and partner rats did differ significantly in emergency latency, irrespective of (early life) background (*MD exp*: test/partner role *F*(1,36) = 15.41, *p* < 0.001, *η^2^* = .30; role*MD *F*(1,36) = 0.30, *p* = .58, *η^2^* = .008 and *CH exp:* test/partner role *F*(1,92) = 9.23, *p* < 0.01, *η^2^* = .09; role*housing *F*(1,92)= 1.88, *p* = .17, *η^2^* = .02). Rats selected as test rats had a shorter latency in these experiments (*MD exp:* test rats mean: 14.6 sec ± 1.5 versus partner rats mean: 39.6 sec ± 6.1, *CH exp:* test rats mean: 17.7 sec ± 2.5 versus partner rats mean: 29.2 sec ± 3.6). Maternal deprivation itself did not affect boldness (MD effect: *F*(1,36) = 0.16, *p* = .69, *η^2^* = .004). Complex housing however did affect boldness (housing effect: *F*(1,92) = 33.39, *p* < 0.001, *η^2^* = .29), with a shorter emergence latency for complex housed animals (standard housed mean: 31.1 sec ± 3.7 versus complex housed mean: 13.6 sec ± 0.9).

### Task development

Three pilot experiments (pilots A-C) were conducted to develop the liberation task. Different liberation configurations were tested in a different order in the three pilots (Figure 1A).

In Pilot A, when working for sucrose in a progressive ratio design, rats earned on average 11.3 ± 1.1 sucrose pellets (Figure 3A, left graph), which roughly corresponds to 40 lever presses to obtain a sucrose pellet. The number of sucrose pellets earned ranged from 8 to 16, meaning some animals stopped at 20 lever presses, while others were willing to perform 118 lever presses for a sucrose pellet. In the liberation task of pilot A (Figure 3A, right graph), *Escape to Box 1, Escape to Box 2* and the *Not Trapped* configurations were tested successively and in this order, and motivation for door opening differed between configurations (Configuration (*F*(2,12) = 11.54, *p* < .01, *η_p_^2^* = .66). Rats completed on average 3.9 ± 1.1 progressive ratios when liberating a cage mate into box 1 (Figure 3A, blue bar). which corresponds to a breakpoint of 7, meaning 7 lever presses are needed to open the door and liberate a cage mate (see figure 2A). The completed ratios ranged from 1 to 9, which means that some animals stopped after the first liberation ratio when a single lever press was needed to open the door, while others continued to press 25 times to obtain a door opening. Liberation into box 2, i.e. liberation without social contact, led to comparable results (post hoc *Escape to Box 1* vs *Escape to Box 2*, *p* = 1.00, Figure 3A orange bar). However, when the partner was not trapped but freely moving in the compartment opposite of the test rat (in box 2, the *Not Trapped* configuration), rats were more motivated to lever press for door opening than in both other configurations (post hoc tests *Escape to Box 1 or 2* vs *Not Trapped, p* < .05 and *p* < .01 respectively; Figure 3A green bar). Rats completed on average 6.8 ± 0.5 progressive ratios, corresponding to about 15 lever presses to obtain door opening.

**Figure 3:**
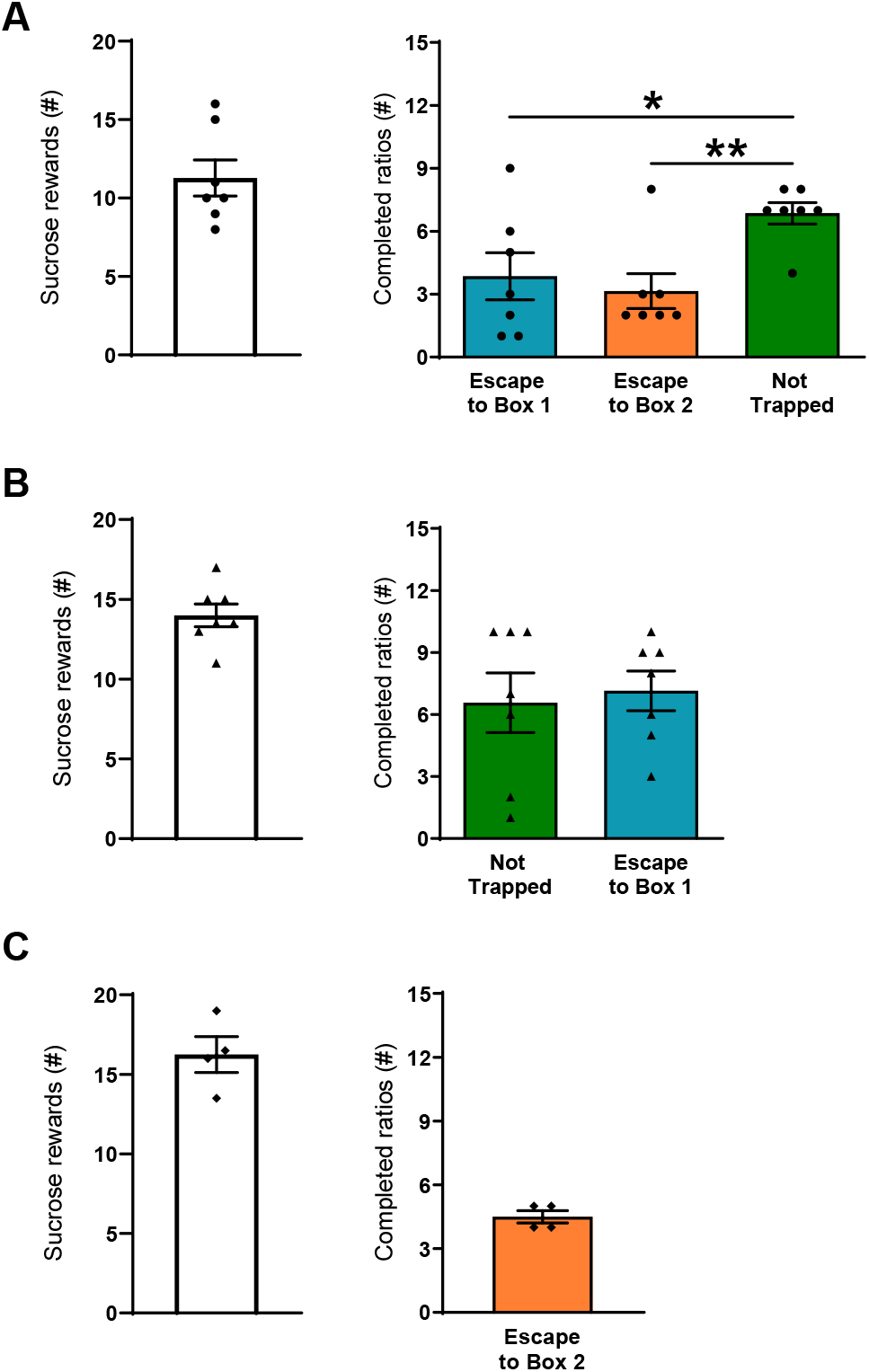
Pilot experiments with the liberation task. The number of sucrose rewards earned in a progressive ratio design is plotted in the graphs on the left (white bars). The motivation to liberate a cage mate (number of completed ratio’s) in an adapted progressive ratio design is plotted in the graphs on the right (blue, orange and green bars). (A) Results for pilot A, where the Escape to box 1 configuration was tested first, followed by Escape to box 2 and the Not Trapped configuration (*n* = 7). (B) Results for pilot B, where the Not Trapped configuration was tested first, followed by Escape to Box 1 (*n* = 7). (C) Results for pilot C, where Escape to Box 2 was tested as a first configuration (*n* = 4). Data represent individual data points and mean ± SEM.

The results of pilot A suggest that test rats show less motivation to open cylinder doors if the partner is trapped compared to when the partner is freely moving. To assess if this difference in motivation was the result of testing order, pilot B and C were conducted. In Pilot B, when working for sucrose, rats earned on average 14 ± 0.7 sucrose pellets (Figure 3B, left graph), corresponding to 77 lever presses to obtain a sucrose pellet. In the liberation task (Figure 3B right graph), when the *Escape to Box 1* configuration was conducted after the *Not Trapped* configuration, there was no difference between the two configurations (*t*(6) = −0.76, *p* = .48, g_av_ = 0.15). The *Not Trapped* configuration in pilot B yielded comparable results to pilot A, but for the *Escape to Box 1* configuration the order of testing mattered. In Pilot C, when working for sucrose, rats earned on average 16.2 ± 1.1 sucrose pellets (Figure 3C, left graph), corresponding to 118 lever presses to obtain a sucrose pellet. Motivation to liberate a cage mate in the *Escape to Box 2* as the first configuration in pilot C (Figure 3C, right graph) was comparable to the results obtained in pilot A. Taken together, this suggests that previous experience is at least important for the *Escape to Box 1* configuration in this task and extensive habituation of the trapped rat to the set-up (during the *Not Trapped* configuration) might account for this. Following these results, the *Escape conditions (to Box 1 or 2)* were always tested as first configurations in the main experiments, before the *not Trapped* condition.

### Maternal deprivation effects on motivation for sucrose and liberation of a trapped cage mate

The motivation to work for a sucrose reward in a progressive ratio design (Figure 4A), was not affected by MD (*t*(18) = 1.02, *p* = .32, g_s_ = 0.44). In the liberation task, MD rats seemed less motivated to liberate a trapped cage mate in the *Escape to Box 1* configuration (Figure 4B), as seen by a lower number of completed ratios, but this effect was (just) not significant, although the effect size was quite large (*t*(18) = 2.05, *p* = .06, g_s_ = 0.88). In the *Not Trapped* configuration (Figure 4C), motivation for door opening was comparable between groups (*t*(18) = 0.22, *p* = .83, g_s_ = 0.09). When tested in this order, rats were more motivated to open the cylinder doors in the *Not Trapped* configuration, compared to the *Escape to Box 1* configuration (Configuration *F*(1,18) = 49.13, *p* < .001, *η_p_^2^* = .73), irrespective of early life experience (Configuration*MD (1,18) = 2.83, *p* = .11, *η_p_^2^* = .14).

**Figure 4:**
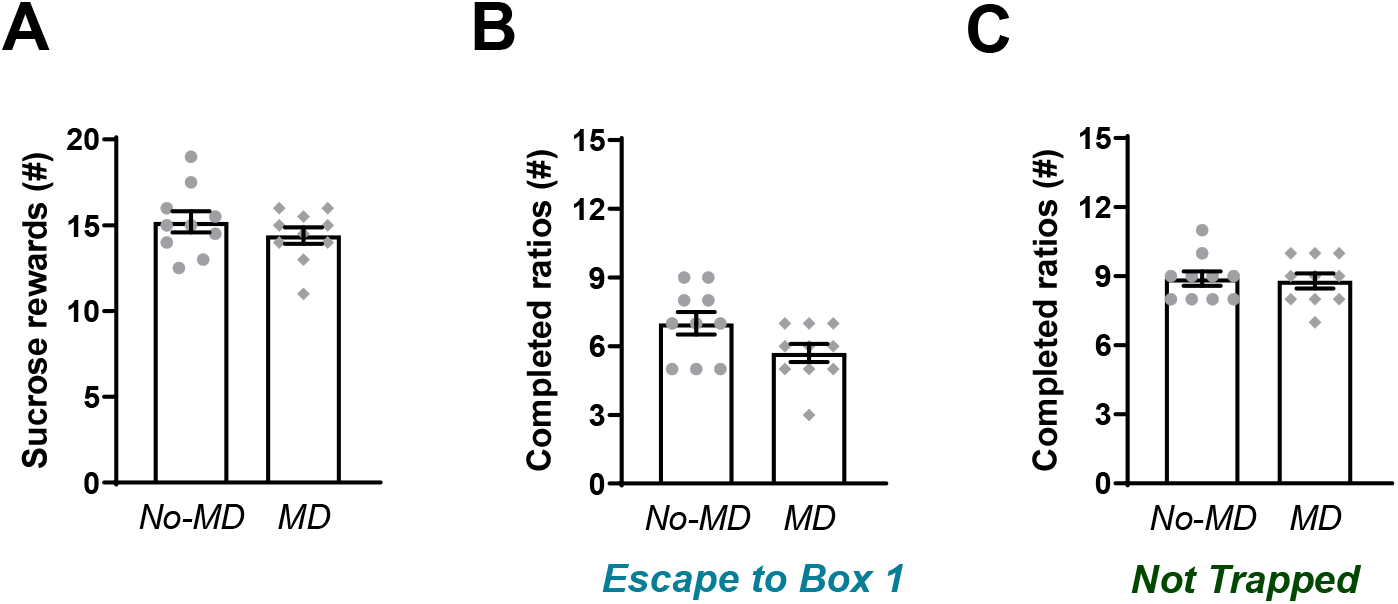
Effect of maternal deprivation on sucrose reward and behavior in the liberation task. (A) The number of sucrose rewards earned in a progressive ratio design. (B) Motivation to liberate a cage mate (number of completed ratio’s) in the *Escape to Box 1* configuration in an adapted progressive ratio design. (C) Motivation to open the doors in the *Not Trapped* configuration. Data represent individual data points and mean ± SEM, *n* = 10 per experimental group.

### Complex housing effects on motivation for sucrose and liberation of a trapped cage mate

All animals were tested on motivation to work for sugar. The number of sucrose rewards earned in a progressive ratio design (Figure 5A and 5B, left graphs) was comparable between standard and complex housed rats, indicating an equal motivation to work for sucrose rewards (1^st^ exp: *t*(46) = −0.29, *p* = .97, g_s_ = 0.00; 2^nd^ exp: *t*(8) = −1.68, *p* = .13, g_s_ = −0.98). When complex housed rats were compared to standard housed rats, they showed a reduced motivation to liberate a trapped cage mate in the *Escape to Box 1* configuration (*t*(46) = 4.13, *p* < .001, g_s_ = 1.15, Figure 5A, middle graph). Similarly, motivation for door opening in the *Not Trapped* configuration was reduced in complex housed rats (*t*(26) = 3.76, *p* < .01, g_s_ = 1.4, Figure 5A right graph). Unlike in earlier experiments, the number of completed ratios did not significantly differ between the Escape to box 1 and Not trapped configurations (Configuration *F*(1,26) = 2.86, *p*= .10, *η_p_^2^* = .09), in either housing condition (Configuration*Housing *F*(1,26) = 0.05, *p* = .83, *η_p_^2^*= .00).

**Figure 5:**
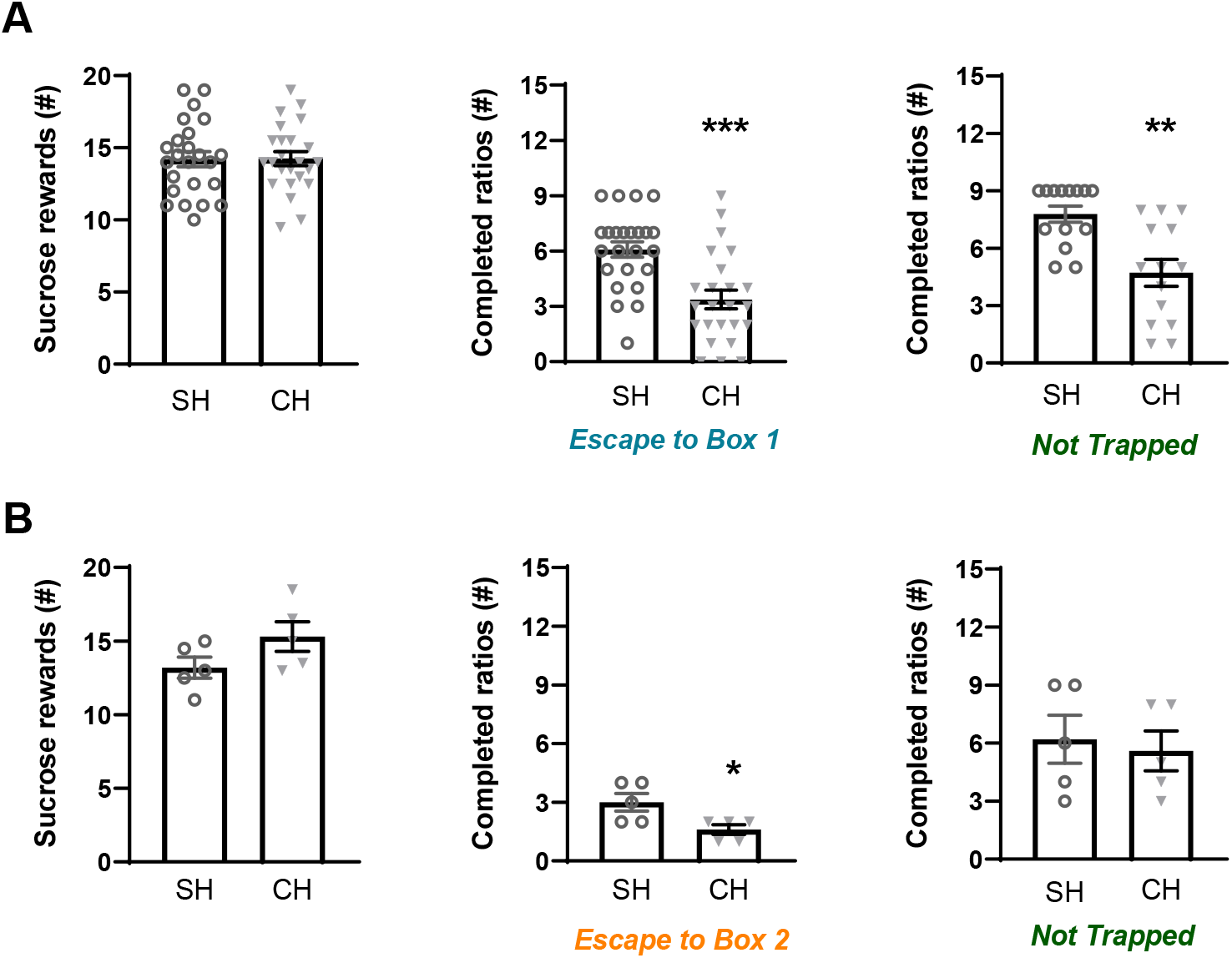
Effect of complex housing on sucrose reward and behavior in the liberation task. The motivation to work for sugar (number of sucrose rewards earned) in a progressive ratio design is depicted in the left graphs. The motivation to liberate a cage mate (number of completed ratios) in an adapted progressive ratio design is depicted in the middle graphs. Motivation to open the doors in the *Not Trapped* configuration is depicted in the right graphs. (A) shows the main experiment with *Sucrose reward (n=24 in each exp group), Escape to box 1 (n=24 in each exp group)* and *not Trapped (n=14 in each exp group)* configurations and (B) shows preliminary data with *Sucrose reward, Escape to box 2 and not Trapped configurations (n=5 in each exp group)*. Data represent individual data points and mean ± SEM.

The preliminary results in the *Escape to box 2* configuration (Figure 5B, middle graph) pointed to a similar direction, that is, complex compared to standard housed rats showed a reduced motivation to liberate a trapped cage mate (*t*(8) = 2.74, *p* < .05, g_s_ = 1.6, Figure 5B middle). This difference was not observed in the *Not Trapped* configuration (*t*(8) = 0.37, *p* = .72, g_s_ = 0.21, Figure 5B right). In this setting, animals were more motivated to lever press for door opening in the *Not Trapped* compared to *Escape to Box 2* configuration (Configuration *F*(1,8) = 14.24, *p* < .01, *η_p_^2^* = .64), irrespective of housing conditions (Configuration*Housing *F*(1,8) = 0.18, *p* = .68, *η_p_^2^* = .02).

When comparing the motivation to work for sugar with the motivation to work for door opening in both housing conditions, we found a correlation between completed ratios in sucrose reward and in the *Escape to Box 1* configuration (SH: *r* = 0.73, *p* < .001; CH: *r* = 0.44, *p* < .05), but not between completed ratios in sucrose reward and the *Not Trapped* configuration (SH: *r* = 0.12, *p*= .68; CH: *r* = 0.49, *p* = .08).

### Ultrasonic vocalizations in the liberation task

In the complex housing experiment, USVs in the first session (FR1) of both the *Escape to Box 1* and *Not Trapped* configuration were recorded and analyzed.

Housing did not affect the emission of alarm calls before door opening (liberation) in either the *Escape to Box 1* configuration (Figure 6A) or the *Not Trapped* configuration (Figure 7A); the emission of 22 kHz calls was comparable between standard and complex housed rats (*Escape to box 1: t*(46) = −1.59, *p* = .12, g_s_ = −0.45 and *Not Trapped: t*(26) = 1.24, *p* = .22, g_s_ = 0.47). The same was true for the emission of appetitive calls in both configurations (Figure 6B/7B); 50 KHz vocalizations before door opening (liberation) were comparable between standard and complex housed rats (*Escape to Box 1: t*(46) = −0.87, *p* = .38, g_s_ = −0.25, *Not Trapped: t*(26) = 1.68, *p* = .10, g_s_ = 0.62).

As a result of door opening (liberation) in the *Escape to Box 1* configuration, USVs emissions significantly decreased for both call types (Figure 6A and 6B), irrespective of housing condition (*alarm calls:* door opening *F*(1,42) = 4.73, *p*<.05, *η_p_^2^*= .10, door opening*housing *F*(1,42) = 2.01, *p*= .16, *η_p_^2^* = .05; *appetitive calls:* door opening *F*(1,42) = 36.33, *p*<.001, *η_p_^2^* = .46, door opening*housing *F*(1,42) = 0.43, *p* = .51, *η_p_^2^* = .01). In the *Not Trapped* configuration (Figure 7A), the emission of alarm calls significantly increased after door opening, irrespective of housing condition (*F*(1,24) = 11.97, *p*<.01, *η_p_^2^* = .33, door opening* housing *F*(1,24) = 1.84, *p* = .19, *η_p_^2^* = .07), although both before and after door opening levels were very low. For the appetitive calls, a significant interaction was found between door opening and housing (*F*(1,24) = 5.87, *p* < .05, *η_p_^2^* = .19), with emission of vocalizations seemingly decreasing after door opening for the standard housed rats and increasing for the complex housed ones. Post hoc analysis however, did not show statistically significant difference between before and after door opening in either group (SH: *t*(13) = 1.6, *p* = .13, g*s* = 0.41, CH: *t*(11) = −1.87, *p* = .09, g_s_ = 0.53).

In general, in both the *Escape to Box 1* and the *Not Trapped* configuration, rats spent more time emitting appetitive calls than alarm calls, both before and after door opening and irrespective of housing conditions (*Escape to box 1:* call type *F*(1,42) = 33.73, *p* < .001, *η_p_^2^* = .44, call type*door opening *F*(1,42) = 1.97, *p* = .05, *η_p_^2^* = .09, call type*housing *F*(1,42) = 0.007, *p*= .93, *η_p_^2^* = .00; *Not Trapped:* call type *F*(1,24) = 46.02, *p* < .001, *η_p_^2^* = .66, call type*door opening *F*(1,24) = 0.25, *p* = .62, *η_p_^2^* = .01, call type*housing *F*(1,24) = 0.60, *p* = .45, *η_p_^2^* = .02).

The emission of alarm calls before liberation and the influence of housing condition could explain a significant amount of the variance observed in the latency to liberate in the *Escape to Box 1*configuration (Figure 6C, *F*(2,45) = 54.59, *p* < .001, *R^2^* = .71); the analysis shows that higher emission rates of alarm calls were observed when liberation of the cage mate took longer (*Beta* = .86, *t*(47) = 10.45, *p* < .001) and complex housed animals tended to have higher liberation latencies (Beta = −.18, *t*(47) = −2.14, *p* < .05). This was not observed for the appetitive calls (Figure 6D, *F*(2,45) = 0.48, *p* = .62, *R^2^* = .02). Thus, in the *Not Trapped* configuration, the emission of alarm calls before door opening and housing condition did not predict the variance observed in the latency to door opening (Figure 7C, *F*(2,25) = 1.06, *p* = .36, *R^2^* = .08), and neither did the appetitive calls (Figure 7D, *F*(2,25) = 0.79, *p* = .46, *R^2^* = .06).

## Discussion

In the present study, we developed an automated, operant pro-social liberation task for rats aimed at measuring motivation to liberate a trapped conspecific. We next investigated the effect of (early) life challenges on performance in this task. Using an adapted version of the pro-social liberation task (Bartal et al., 2011), rats learned to press a lever that opened doors of a cylinder in which a partner rat was trapped. With this task we showed that rats are motivated to liberate a trapped cage mate, even when liberation does not lead to social contact. Moreover, we showed that rats are more motivated to lever press and gain access to the cage mate when the cage mate is not trapped but freely moving in a separate compartment. This suggests that rats are affected by the emotional state of the partner and adapt their behavior depending on whether or not the partner is trapped, though in this case trapping of the partner *reduced* motivation to act. In support of this finding, analysis of USVs revealed that there was a positive correlation between the amount of 22 kHz alarm calls (and not 50KHz appetitive calls) emitted before liberation and the latency to door opening, suggesting that increased levels of distress were linked to a decreased motivation for door opening. Manipulation of the early postnatal environment by 24h MD at postnatal day 3 did not affect behavior in this task, regardless of whether the cage mate was trapped or freely moving in a separate compartment. Rearing rats in complex housing from postnatal day 26 onwards did affect behavior; compared to standard housed rats, complex housed rats displayed a similar motivation to press a lever for sucrose, yet a reduced motivation to liberate a trapped cage mate as well as a reduced motivation to gain access to a non-trapped rat. During the first test session with either a trapped or not trapped cage mate, emission of both 22kHz and 50kHz calls was comparable between standard housed and complex housed rats, suggesting that USVs are not necessarily linked to the reduced motivation of door opening in the complex housed animals.

### Task development

Our paradigm was inspired by the seminal work of Bartal and colleagues (Bartal et al., 2011, 2014, 2016). In these studies rats learned over time to manually open a cylinder that contains a trapped conspecific. As the requirements for door opening do not change over the course of testing, an increase in the percentage of door openings and decrease in door opening latencies over sessions demonstrated that liberation of the cage mate is reinforcing and stimulates repeated liberation. A similar study with similar findings was performed by Sato et al. (2015), in which the partner was trapped in a pool of water instead of a cylinder.

In our set-up the requirements for door opening increased over sessions –to probe motivation-until the rats ceased to meet the requirements and the experiment ended. Because we measured motivation in a progressive ratio design, this allowed us to compare motivation to work for a social reward with motivation to work for a food reward. Similar to the current study, Bartal and colleagues separated the desire for social contact from ending the distress of a conspecific in a control experiment in which the rat was separated into a second compartment and showed that the desire for social contact was not the main force driving behavior. The current data set complements these earlier studies, by replicating the finding that rats are willing to liberate a trapped conspecific repeatedly and with increased cost. Moreover, we introduced a non-trapped configuration, and show that the motivation to gain access to a freely moving cage mate is higher than the motivation to liberate a trapped cage mate with subsequent social contact.

More specifically, three different configurations were tested to distinguish between motivation to work for social contact and motivation for liberating a conspecific from an aversive situation. In the *Escape to Box 1* configuration the trapped partner is liberated into the test rat compartment, which means that motivation to liberate is potentially driven by a mixture of the desire for social contact and liberating the conspecific from an aversive situation. In order to study the weight of these factors separately, the *Not Trapped and Escape to Box 2* configurations were tested as well. Eliminating the element of social contact in the *Escape to Box 2* configuration did not change the number of completed ratios accomplished, neither when tested as a second configuration, nor when tested as the starting configuration. Although this does not prove that the desire for social contact does not play a role in the *Escape to Box 1* configuration, it does not seem to be the main driving force. Next, the *Not Trapped* configuration was tested in which the partner was freely moving in a second compartment that was separated from the test rat by the cylinder and closed doors. Test rats could lever press to open the doors and gain access to the partner in the second compartment. Rats were more motivated to work for door opening in this setting than in configurations where the partner was trapped, both when tested in sequence to the trapped configuration and when tested as a first configuration. The finding that liberation of the trapped rat after testing the *Not Trapped* configuration yielded comparable result between both settings (pilot B, Figure 3C), might be due to the fact that the partner had ample opportunity to voluntarily explore the cylinder and habituate to it (with its cage mate present), rendering the following “trapped” sessions less stressful. Future recordings of USV during these sessions might support this interpretation.

If, through emotional contagion, rats are negatively affected by the distressed state of the trapped partner, rats are expected to downregulate their own distress by liberating the partner. When we started testing, we placed the partners in a much smaller cylinder in which animals could not move or turn and showed clear physical signs of distress (such as actively struggling, excretion of feces and urinating). In this setting, test rats did not press the lever at all, but rather avoided the cylinder. This avoidance of stressed conspecifics by male rats has also been reported by others (Rogers-Carter et al., 2018). On the other hand, Bartal and colleagues (2016) indicated that rats that were less stressed were also less inclined to show pro-social behavior. In the current setting, animals did press the lever to liberate a cage mate, but the number of completed ratios mostly remained lower than in the *Not Trapped* configuration. Taken together, data from the current and other studies show that there is a delicate window in which vicarious distress induces other rats to act pro-socially.

Both 22kHz alarm calls and 50kHz appetitive calls were recorded during the first test session (FR1) of both the Trapped and Not Trapped configurations. If liberation of a trapped rat is motivated by a shared affective state via emotional contagion, then ultrasonic communication is a prime candidate for the transfer of emotions between individuals (Brudzynski, 2013). In the Trapped configuration, the amount (depicted in percentage of time that calls were emitted) of both 22kHz alarm and 50KHz appetitive calls decreased after door opening, which was not observed in the Not Trapped configuration. This indicates that USVs may play a larger role in communication when the partner is trapped and subsequently liberated than in non-trapped conditions. It is to be noted that USVs were recorded per pair of animals and it was not possible to identify the source of the alarm calls (test rat or partner). Ultrasonic communication before liberation also correlated with the behavior of the test rat; i.e. higher emission rates of 22 kHz alarm calls coincided with longer door opening latencies. In light of our previous observations we speculate that higher emission rates of 22kHz calls delay door opening latency, but it might also be true that longer door opening latencies increase 22kHz vocalization. Nonetheless, the positive correlation between alarm calls and liberation latency suggests that ultrasonic communication might contribute to the process of emotional contagion and that increased levels of distress might reduce active helping behavior. These preliminary results clearly await more in-depth investigation.

### Challenging early life conditions

Two separate experimental conditions were introduced to study how the (early) life environment affects development of pro-social behavior. First, the early postnatal environment was manipulated by depriving pups on postnatal day 3 from maternal care for 24h. In rodents, stressful experiences in the early postnatal period, including maternal deprivation, can alter stress reactivity and cause deficits in cognitive and emotional behavior (Lupien et al., 2009; Krugers and Joëls, 2014; Marco et al., 2015; Van Bodegom et al., 2017; Bonapersona et al., 2019). Regarding social behavior, early life stress studies have found limited effects on social interest, but mostly a decrease in social discrimination and a decrease in (or no effect on) social interaction (van der Veen et al., 2020). Whether or not an animal acts to help a distressed conspecific is a complex context-dependent process and it is likely that individual behavioral characteristics such as anxiousness and ability to self-regulate emotions affect the response to distressed others.

To our knowledge, the current research is the first to study effects of early life stress on pro-social behavior. Compared to non-deprived controls, MD animals tended to show lower levels of pro-social activity in the Trapped configuration, with a quite high effect size, but not reaching statistical significance, most likely due to the relatively small sample sizes. Performance in the Not Trapped configuration was very similar in MD rats and non-deprived controls. As part of lever acquisition training, rats also conducted a progressive ratio for sucrose which allowed us to compare motivation to work for a food reward with motivation to work for a social reward. MD animals were equally motivated to work for a sucrose reward as non-deprived controls. Together with performance in the liberation task, it seems that reward sensitivity, both for food and a social reward, is not strongly affected by MD, at least not in the current experimental design.

A second environmental manipulation concerned complex housing from early adolescence onwards, starting on postnatal day 26 and lasting until testing. Complex housed rats were less motivated to liberate a trapped cage mate, as reflected by a significant reduction in the number of completed ratios compared to standard housed controls in both the *Escape to box 1* (towards) and *Escape to box 2* (away from) configurations. This reduction in motivation is probably not explained by increased emotional contagion or reduced behavioral self-regulation, because motivation in the *Not Trapped* configuration was also reduced. Instead, reduced motivation to act in both configurations might reflect an overall reduced motivation to work for a social reward. Moreover, this decrease in motivation appeared to be specific for social rewards, as the motivation to work for a sucrose reward was comparable for complex and standard housed rats. These results seem to be in accordance with recent findings of our lab where complex housed rats were tested in a pro-social two-choice task. In this task, rats were given the choice between lever pressing to obtain a sucrose reward for themselves or for themselves and their partner. Standard housed rats on average preferred the pro-social option, while complex housed rats did not (Kentrop et al., 2020). We have also previously shown that testing complex housed rats shortly after taking them from the (complex) home cage reveals a strong reduction in social interest compared to standard housed animals (Kentrop et al., 2018). This reduced social interest -as a result of living with 10 conspecifics-could also affect the reward experienced by liberating or gaining access to another rat and lead to less pro-social behavior.

Previous experiments have shown that complex housed rats quickly adjust to novel environments and show a reduced behavioral response to mild novelty stress (Zimmermann et al., 2001; van der Veen et al., 2015; Kentrop et al., 2018). We hypothesized that complex housed partners might experience less distress from being trapped in the cylinder than standard housed partners, which, as a result, could lower the urgency for complex housed rats to liberate their partner. However, the emission of 22 kHz alarm calls before liberation was equal in complex and standard housed groups, suggesting that at least in the first test session partners from both groups experienced equal distress from being trapped. Interestingly, the emission of 50 kHz calls was also higher before than after door opening. These 50 kHz calls however were not correlated with liberation latency. This pattern of results suggests more communication in general before liberation.

In conclusion, we developed and tested an adapted version of a behavioral task that measures pro-social behavior in rats. We show suggestive evidence that rats are sensitive to the distressed state of trapped cage mates and are willing to liberate them; and that this might be affected by challenging conditions during (early) life. Moreover, the task demonstrated to be sufficiently sensitive to measure motivation for door opening in different contexts, which allows for future experiments that examine contextual prerequisites needed for individuals to behave pro-socially.

## Acknowledgments

We thank Valeria Bonapersona, Jos Brits, Ruth Damsteegt and Jelle Knop for their assistance during experimental procedures. This work was supported by *the Consortium on Individual Development (CID)*, which is funded through the Gravitation program of the Dutch Ministry of Education, Culture, and Science and the Netherlands Organization for Scientific Research (NWO grant number 024.001.003). M.J. Bakermans-Kranenburg was supported by the European Research Council (ERC AdG 669249); M.H. van IJzendoorn was supported by the Netherlands Organization for Scientific Research (Spinoza Prize). The authors declare that the research was conducted in the absence of any professional or financial relationships that could be construed as a potential conflict of interest.

